# Protein Docking Model Evaluation by Graph Neural Networks

**DOI:** 10.1101/2020.12.30.424859

**Authors:** Xiao Wang, Sean T Flannery, Daisuke Kihara

## Abstract

Physical interactions of proteins play key roles in many important cellular processes. Therefore, it is crucial to determine the structure of protein complexes to understand molecular mechanisms of interactions. To complement experimental approaches, which usually take a considerable amount of time and resources, various computational methods have been developed to predict the structures of protein complexes. In computational modeling, one of the challenges is to identify near-native structures from a large pool of generated models. Here, we developed a deep learning-based approach named Graph Neural Network-based DOcking decoy eValuation scorE (GNN-DOVE). To evaluate a protein docking model, GNN-DOVE extracts the interface area and represents it as a graph. The chemical properties of atoms and the inter-atom distances are used as features of nodes and edges in the graph. GNN-DOVE was trained and validated on docking models in the Dockground database. GNN-DOVE performed better than existing methods including DOVE, which is our previous development that uses convolutional neural network on voxelized structure models.

## 1 Introduction

Experimentally determined protein structures provide fundamental information about the physicochemical nature of biological function of protein complexes. With the recent advances in cryo-electron microscopy, the number of experimentally determined protein complex structures has been increasing rapidly. However, experimental methods are costly in terms of money and time. To aid the experimental efforts, computational modeling approaches for protein complex structures, often referred to as protein docking (*1*), have been extensively studied over the past two decades.

Protein docking methods aim to build the overall quaternary structure of a protein complex from the tertiary structure information of individual chains. Similar to other protein structure modeling methods, protein docking can also be divided into two main categories: template-based methods (*2, 3*), which use a known structure as a scaffold of modeling, and *ab initio* methods, which assemble individual structures and score generated models to choose most plausible ones. In *ab initio* methods, various approaches were used for molecular structure representations (*4*) (*5*). These include docking conformational searches, such as Fast Fourier Transform (*6, 7*), geometric hashing (*4, 8*), and particle swarm optimization (*9*), and for considering protein flexibility (*10, 11*). Development of new methods aim to extend and surpass the capabilities of simple pairwise docking, such as multi-chain docking (*12-14*), peptide-protein docking (*15-17*), docking with disordered proteins (*18*), docking order prediction (*19, 20*), and docking for cryo-EM maps (*21, 22*). Researchers have also applied recent advances in deep learning to further boost docking performance (*23-25*).

Although substantial improvements have been made in *ab initio* protein docking, selecting near-native (i.e. correct) models out of a large number of produced models, which are often called decoys, is still challenging. The difficulty is partly due to a substantial imbalance of the number of near-native models and incorrect decoys in a generated decoy pool. The accuracy of scoring decoys certainly determines the overall performance of protein docking, and thus there are active developments of scoring functions (*26*) for docking models. Recognizing the importance of scoring, the Critical Assessment of PRediction of Interactions (CAPRI) (*27*), which is the community-based protein docking prediction experiments, has arranged a specific category of evaluating scoring methods, where participants are asked to select 10 plausible decoys from over thousands of decoys provided by the organizers. Over the last two decades, various approaches have been developed for scoring decoys. The main categories include physics-based potentials (*11*), scoring based on interface shape (*4, 28*), knowledge-based statistical potentials (*29, 30*), machine learning methods (*31*), evolutionary profiles of interface residues (*32*), and deep learning methods using interface structures (*33*).

In our previous work, we developed a model selection method for protein docking, DOVE (*33*), which uses Convolutional deep Neural Network (CNN) as the core of its architecture. DOVE captures atoms and interaction energies of atoms located at the interface of a docking model using a cube of 20^3^ or 40^3^ Å^3^ and judges if the model is correct or incorrect according to the CAPRI criteria (*34*). We showed that DOVE performed better than existing methods. However, DOVE has a critical limitation - since it captures an interface with a fixed-size cube, only a part of the interface is captured when the interface region is too large. This often caused an erroneous prediction. In addition, a 3D grid representation of an interface often includes voxels of void space where no atoms exist inside, which is not efficient in memory usage and may even be detrimental for accurate prediction. In this work, we address this limitation of DOVE by applying a graph neural network (GNN)(*35, 36*), which has previously been successful in representing molecular properties (*37-40*). Using GNN allows the capturing of all interface atoms in a more flexible and natural fashion. To the best of our knowledge, this is the first method that applies GNNs to the protein docking problem. Compared to DOVE and other existing methods, GNN-DOVE demonstrated substantial improvement in a benchmark study.

## 2 Materials and methods

We first introduce the datasets used for training and testing GNN-DOVE. Subsequently, we introduce the graph neural network architecture and the training process of GNN-DOVE.

### 2.1 Dataset

To train and test GNN-DOVE, we used the Dockground dataset (*41*). The dataset includes 58 target complexes, each with on average 9.83 correct and 98.5 incorrect decoys. Correct decoys were identified by applying the CAPRI criteria (*27*), which considers interface root mean square deviation (iRMSD), ligand RMSD (lRMSD), and the fraction of native contacts (fnat). The iRMSD is the Cα RMSD of interface residues with respect to the native structure. Interface residues in a complex are defined as all the residues within 10.0 Å from any residues of the other subunit. lRMSD is the Cα RMSD of ligands when receptors are superimposed, and fnat is the fraction of contacting residue pairs, i.e. residue pairs with any heavy atom pairs within 5.0 Å, that exist in the native structure.

To remove redundancy, we grouped the 58 complexes using the TM-Score (*42*). Two complexes were assigned to the same group if at least one pair of proteins from the two complexes had a TM-score of over 0.5 and sequence identity of 30% or higher. This resulted in 29 groups (Table 1). In Table 1, complexes (PDB IDs) of the same group are shown in lower case in a parenthesis followed by the PDB ID of the representative. These groups were split into four subgroups to perform four-fold cross-testing, where three subsets were used for the training while one subset was used for testing. Thus, by cross-testing, we have four models tested on four fully independent testing sets. Among the training set, we used 80% of decoys for training a model and the remaining 20% as a validation set, which was used to determine the best hyper-parameter set for training. In the results, the accuracy of targets when treated in the testing set was reported. To have a fair comparison with DOVE (*33*), DOVE was also trained and tested using this protocol.

**Table 1.** Dataset splits for training and testing GNN.

| Fold | PDB ID |
| --- | --- |
| 1 | 1A2K, 1E96 (1he1, 1he8, 1wq1), 1F6M, 1MA9(2btf), 1G20, 1KU6, 1T6G, 1UGH, 1YVB, 2CKH, 3PRO |
| 2 | 1AKJ (1p7q, 2bnq), 1DFJ, 1NBF (1r4m, 1xd3, 2bkr), 1GPW, 1HXY, 1U7F, 1UEX, 1ZY8, 2GOO, 1EWY |
| 3 | 1AVW (1bth, 1bui, 1cho, 1ezu, 1ook, 1oph, 1ppf, 1tx6, 1xx9, 2fi4, 2kai, 1r0r, 2sni, 3sic) |
| 4 | 1BVN (1tmq), 1F51, 1FM9, 1A2Y (1g6v, 1gpq, 1jps, 1wej, 1l9b, 1s6v), 1W1I, 2A5T, 3FAP |

There are in total 29 representative targets shown in the upper case, targets in the lower case in a parenthesis indicate that they belong to the same group.

### 2.2 The GNN-DOVE algorithm

In this section, we describe GNN-DOVE, which uses the graph neural network. The GNN-DOVE algorithm is inspired by a recent work in drug-target interactions (*38*), which designed two graph-representation for capturing intermolecular interactions for protein-ligand interactions. We will first explain how the 3D structural information of a protein-complex interface is embedded as a graph. Then, we describe how we used a graph attention mechanism to focus on the intermolecular interaction between a receptor and a ligand protein. The overall protocol is illustrated in Figure 1. For an input protein docking decoy, the interface region is identified as a set of residues located within 10.0 Å of any residues of the other protein. A residue-residue distance is defined as the shortest distance among any heavy atom pairs across the two residues. Using the extracted interface region, two graphs are built representing two types of interactions: the graph *G*^1^ describes heavy atoms at the interface region, which only considers the covalent bonds between atoms of interface residues within each subunit as edges. Another graph *G*^2^ connects both covalent (thus includes *G*^1^ and non-covalent residue interaction as edges, where a non-covalent atom pair is defined as those which are closer than 10.0 Å from each other. Both graphs will be processed by graph neural network (GNN) to output a score, which is a probability that the docking decoy has a CAPRI acceptable quality (thus making higher scores better).

**Figure 1.**
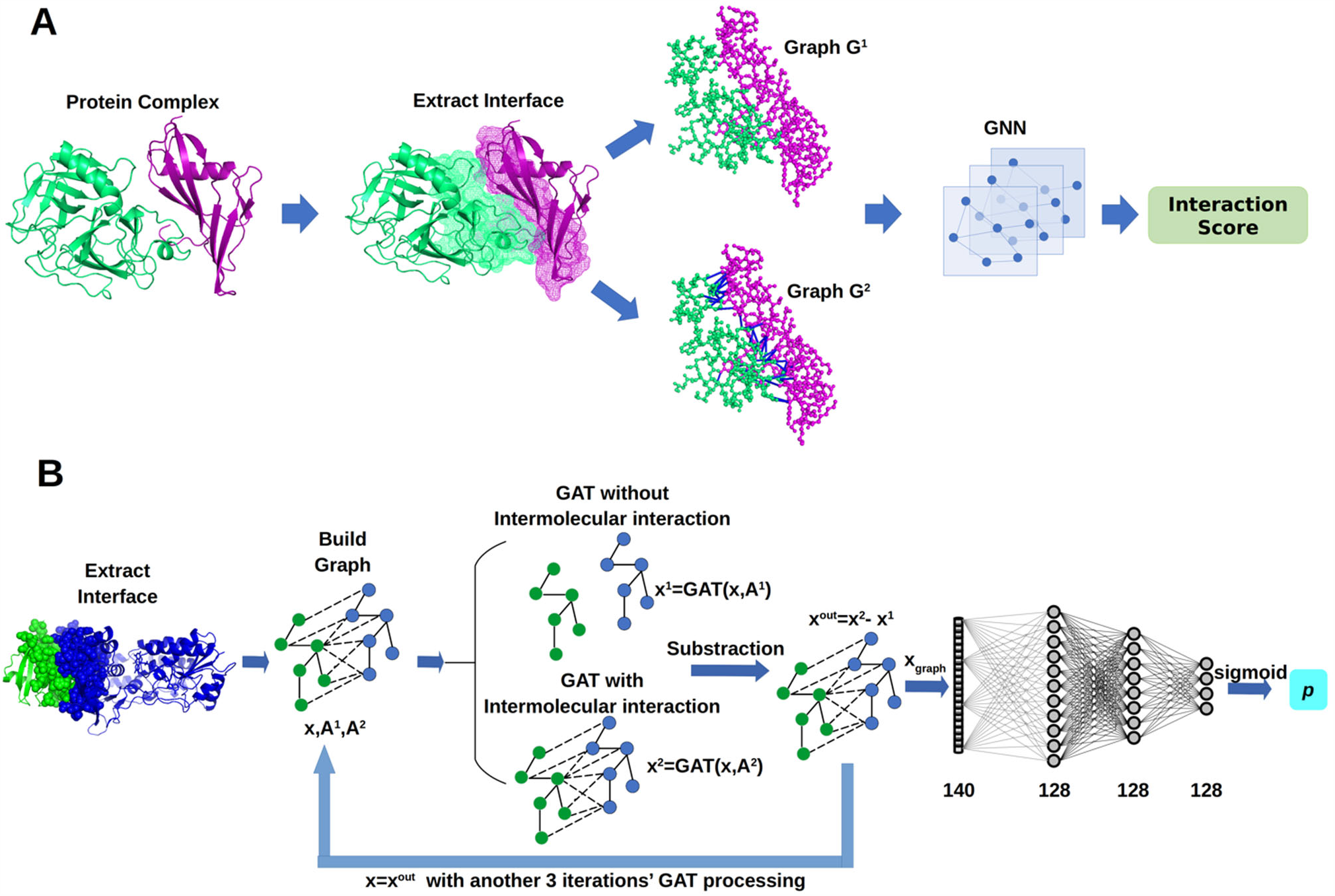
The framework of GNN-DOVE. GNN-DOVE extracts the interface region of protein complex and further reconstructs graph with/without intermolecular interactions as input, then outputs the probability that indicates if the input structure is acceptable or not. **A**, the overall logical steps of the pipeline. **B**, the architecture of the GNN network with the GAT mechanism.

#### Building Graphs

A key feature of this work is the graph representation of an interface region of a complex model. Graph *G* is defined by *G = (V, E, A)*, where *V* denotes the node set, *E* is a set of edges, and *A* is the adjacency matrix, which numerically represents the connectivity of the graph. For a graph *G* with *N* nodes, the adjacency matrix *A* has a dimension of *N*N*, where *A*_*ij*_ > 0 if the *i-*th node and the *j-*th node are connected and *A*_*ij*_ *=* 0 otherwise. The adjacency matrix *A*^1^ for graph *G*^1^ describes covalent bonds at the interface, and thus defined as follows:

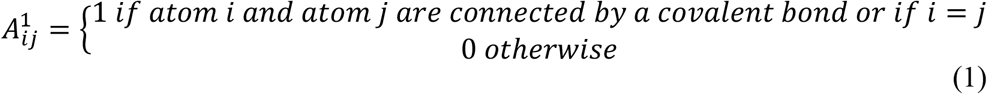

The matrix *A*^2^ for *G*^2^ describes both covalent bonds and non-covalent interactions between atoms within 10.0 Å to each other. It is defined as follows:

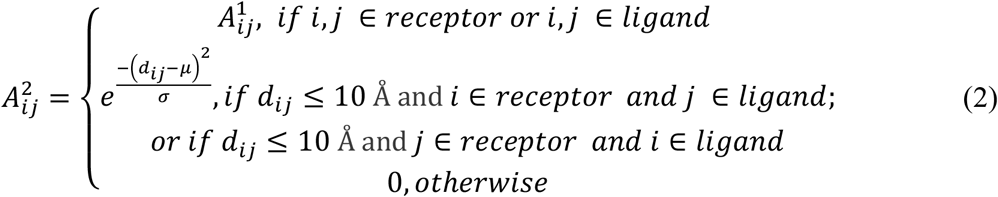

where *d*_*ij*_ denotes the distance between the *i-*th and the *j-*th atoms. *μ* and *σ* are learnable parameters, whose initial value is 0.0 and 1.0. The formula 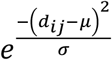 decays as the distance increases between atoms.

Compared to the previous voxel representation used in DOVE, the graph representation encodes the distance information more flexibly and naturally. Note that the representation is rotationally invariant. Also, memory usage is more efficient as void spaces are not represented as is needed for the voxel representation.

As for the node features in the graph, we considered the physicochemical properties of atoms. We used the same features as used in previous works (*38, 43*) as shown in Table 2.

**Table 2.**
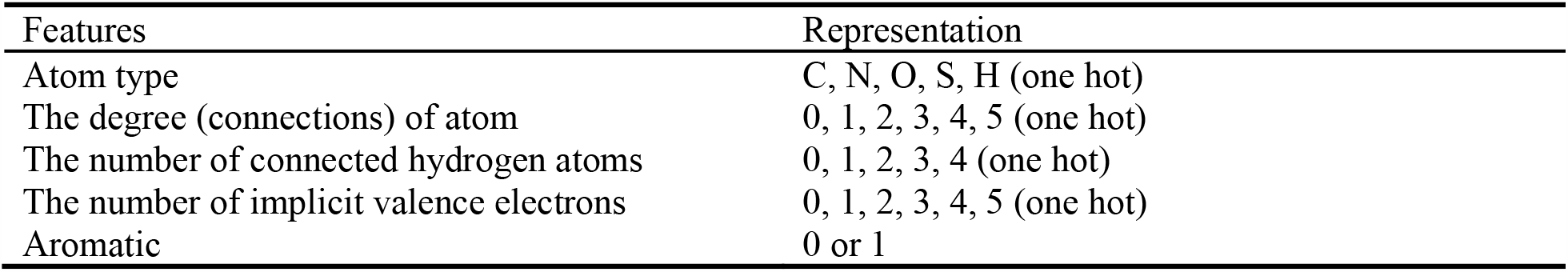
Atom Features

| Features | Representation |
| --- | --- |
| Atom type | C, N, O, S, H (one hot) |
| The degree (connections) of atom | 0, 1, 2, 3, 4, 5 (one hot) |
| The number of connected hydrogen atoms | 0, 1, 2, 3, 4 (one hot) |
| The number of implicit valence electrons | 0, 1, 2, 3, 4, 5 (one hot) |
| Aromatic | 0 or 1 |

#### Attention and Gate-Augmented Mechanism

The constructed graphs are used as the input to the GNN. More formally, graphs are the adjacency matrix *A*^1^ and *A*^2^, and the node features, 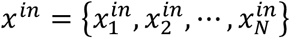 with *x* ∈ ℝ^*F*^, where *F* is the dimension of the node feature.

We first explain the attention mechanism of GNN. With the input graph of *x*^*i*^, the pure graph attention coefficient is defined in Eq. (3), which denotes the relative importance between the *i-*th and the *j-*th node:

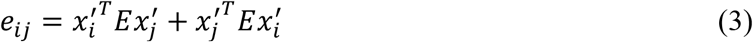

where 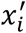 and 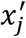 is the transformed feature representation defined by 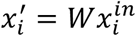 and 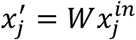. *W, E* ∈ ℝ^*F*×*F*^ are learnable matrices in GNN. *e*_*ij*_ and *e*_*ji*_ become identical to satisfy the symmetrical property of the graph by adding 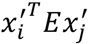 and 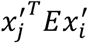. The coefficient will only be computed for *i* and *j* where *A*_*ij*_ > 0.

Attention coefficients will be also computed for elements in the adjacency matrices. They are formulated in the following form for the element (*i, j*):

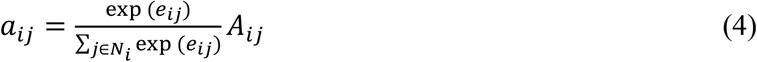

where *a*_*ij*_ is the normalized attention coefficient for the *i-*th and the *j-*th node pair, *e*_*ij*_ is the symmetrical graph attention coefficient computed in Eq. (3), *N*_*i*_ is the set of neighbors of the *i-*th node that includes interacting nodes *j* where *A*_*ij*_ > 0.

Based on the attention mechanism, the new node feature of each node is updated by considering its neighboring nodes, which is a linear combination of the neighboring node features with the final attention coefficient *a*_*ij*_:

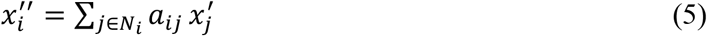

Furthermore, the gate mechanism is further applied to update the node feature since it is known to significantly boost the performance of GNN. The basic idea is similar to that of ResNet (*44*), where the residual connection from the input helps to avoid information loss, alleviating the gradient collapse problem of the conventional backpropagation. The gated graph attention can be viewed as a linear combination of 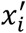 and 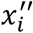, as defined in Eq. (6):

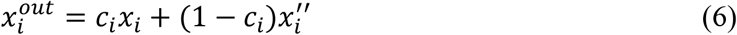

where 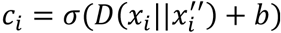, where *D* ∈ ℝ^2*F*^ is a weight vector that is multiplied (dot product) with the vector 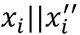 and *b* is a constant value. Both D and b are learnable parameters and share among different nodes. 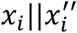 denote the concatenation vector of *x*_*i*_ *and* 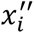.

We refer the Attention and Gate-Augmented Mechanism as a layer, the gate augmented graph attention layer (GAT). Then, we can simply denote 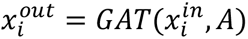. The node embedding can be iteratively updated by *GAT*, which aggregates information from neighboring nodes.

#### Graph Neural Network Architecture of GNN-DOVE

Using the *GAT* mechanism described before, we adopted four layers of *GAT* in GNN-DOVE to process the node embedding information from neighbors and to output the updated node embedding (Figure 1B). For the two adjacency matrices *A*^1^ and *A*^2^, we used the shared (i.e. the same) GAT. The initial input of the network is atom features. With two matrices, *A*^1^ and *A*^2^, we have *x*_1_ *= GAT(x*^*in*^, *A*^1^) and *x*_2_ *= GAT(x*^*in*^, *A*^2^). To focus only on the intermolecular interactions within an input protein complex model, we subtracted the embedding of the two graphs as the final node embedding. By subtracting the updated embedding *x*_1_ from *x*_2_, we can capture the aggregation information that only comes from the intermolecular interactions with other nodes in the protein complex model. Thus, the output node feature is defined as:

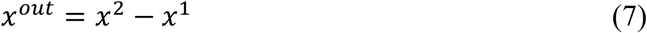

Then, the updated *x*^*out*^ will become *x*^*in*^ to iteratively augmented the information through three following *GAT* layers. After the node embeddings were updated by the four *GAT* layers, the node embedding of the whole graph was summed up as the entire graph representation, which is considered as the overall intermolecular interaction representation of the protein complex model:

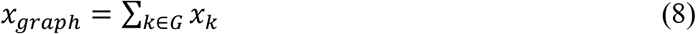

Finally, fully connected (FC) layers were applied to *x*_*graph*_ to classify whether the protein-complex model is correct or incorrect. In total, four FC layers were applied. RELU activation functions were used between the FC layers, and a sigmoid function was applied for the last layer to output a probability value.

The source code of GNN-DOVE is available at https://github.com/kiharalab/GNN_DOVE.

### 2.3 Training nets

Since the dataset was highly imbalanced with more incorrect decoys than acceptable ones, we balanced the training data by sampling the same number of acceptable and incorrect decoys in each batch for training.

The dimension of the graph attention layer (GAT) is 140, while the fully connected layer (FC) is 128. For training, cross-entropy loss (*45*) was used as the loss function optimize the GNN. The Adam optimizer (*46*) with a learning rate of 0.002 was used. Each model of different folds was trained for 100 epochs with a batch size of 32. To avoid overfitting, a dropout (*47*) of 0.3 was applied for every layer except the last FC layer. Weights of every layer were initialized using the Glorot-uniform (*48*) to have a zero-centered Gaussian distribution, and bias was initialized to 0 for all layers. For different fold training, we adopted the same hyper-parameters. The training process generally converged after approximately 30 epochs.

### 2.4 DOVE

We compared the performance of GNN-DOVE with its predecessor, DOVE. Here we briefly describe the DOVE algorithm. DOVE is a CNN-based method for evaluating protein docking models. It first extracts the interface region of an input protein complex model, and the region is put into a 40*40*40 Å^3^ cube as input. A seven-layer CNN, which consists of three convolutional layers, two pooling layers, and two fully connected layers, was adopted to process the voxel input. The output of DOVE is the probability that indicates whether the input model is acceptable or not. For input features, DOVE took atom types as well as atom-based interaction energy values from GOAP (*49*) and ITScore (*29*). Since voxelized structure input is not rotation invariant, DOVE needed to augment training data by rotations.

## 3 Results

### 3.1 Performance on the Dockground dataset

We evaluated the performance of GNN-DOVE on the Dockground dataset. GNN-DOVE was compared with DOVE and five other existing structure model scoring methods, GOAP (*49*), ITScore (*29*), ZRANK (*50*), ZRANK2 (*51*), and IRAD (*52*). The test set results were reported for GNN-DOVE and DOVE. Both GOAP and ITScore were run in two different ways. First, as originally designed, the entire complex structure model was input. The other way was to input only the interface residues that are within 10 Å from the interacting protein (denoted as GOAP-Interface, ITScore-Interface). Thus, GNN-DOVE was compared with a total of eight methods. As for DOVE, we used DOVE with a cube size of 40^3^ Å^3^ and heavy atom distribution as input feature because this setting performed the best among other settings tested on the Dockground dataset in the original paper (*53*) (Figure 4 in the paper, the setting was named as DOVE-Atom40). For this work, DOVE was newly retrained using the same four-fold cross-testing as GNN-DOVE.

Figure 2 shows the hit rate of GNN-DOVE in comparison with the other methods. A hit rate of a method is the fraction of target complexes where the method ranked at least one acceptable model based on the CAPRI criteria within each top rank. In Figure 2, we show two panels. Panel A shows the fraction of targets where a method had at least one hit among each rank cutoff, while in Panel B, the hit rates for a method were averaged first within each of the 29 groups then re-averaged over the groups.

**Figure 2.**
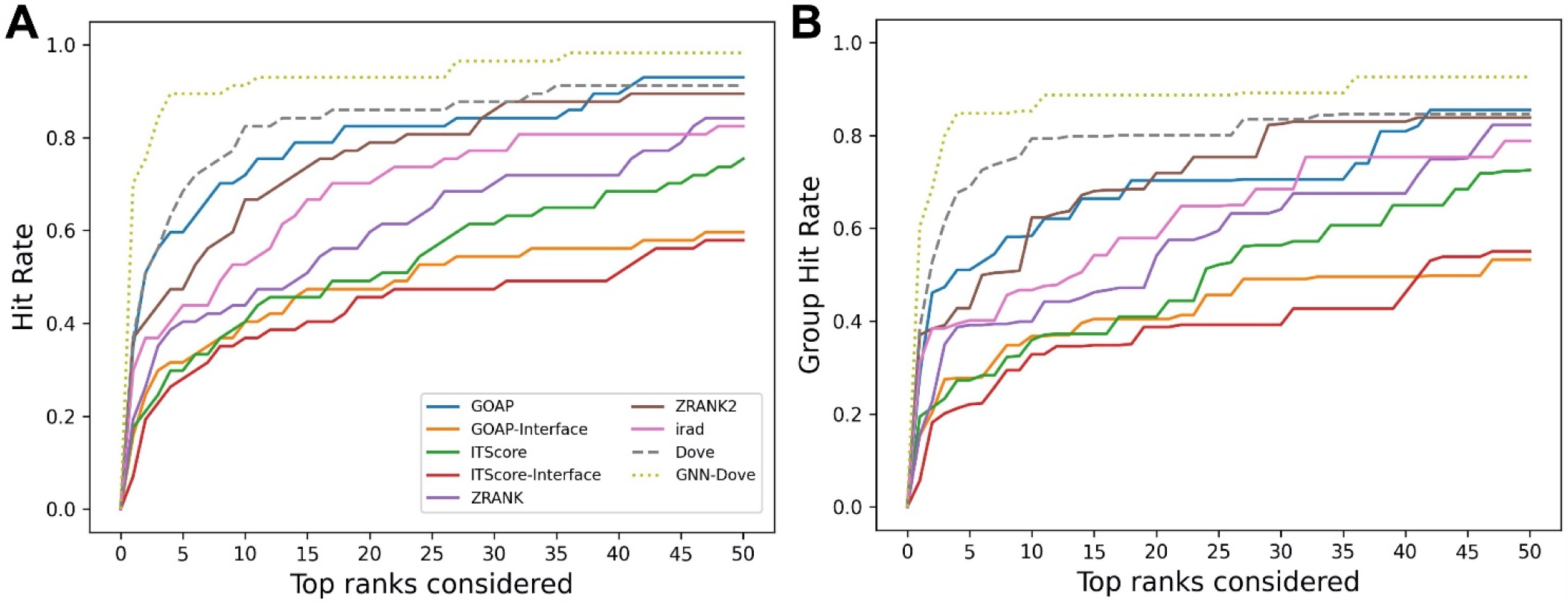
The performance on the Dockground dataset. GNN-DOVE was compared with DOVE and seven other scoring methods. **A**, The panel shows the fraction of target complexes among the 58 complexes in the benchmark set for which a method selected at least 1 acceptable model (within top *x* scored models). **B**, Considering the complexes are grouped into 29 groups, we also compared the hit rate of different methods based on the group classification. The hit rates for complexes in each group were averaged and then re-averaged over the 29 groups.

It is clear from Figure 2 that GNN-DOVE (dotted line in light green) performed better than the other methods. GNN-DOVE was able to rank correct models within earlier ranks in many target complexes. Within the top 10 rank, GNN-DOVE achieved a hit rate of 89.7% while the next best method, DOVE, achieved 81.0% and the third best method, GOAP, obtained 70.7% (Figure 2A). When we further compared the hit rates considering the target groups (Figure 2B), GNN-DOVE consistently outperformed other methods. The gap between GNN-DOVE and DOVE against the other existing methods also increased. Among the other seven existing methods, GOAP showed the highest hit rate at 5^th^ ranks followed by Zrank2 in both panels while ITScore-Interface had the lowest hit rates on this dataset.

In Figure 3, we compared iRMSD, lRMSD, and fnat values of the methods. These metrics are used for defining the quality levels in CAPRI. The best value among the top 10 ranked decoys was plotted. For the majority of the cases (49 out of the 58 targets) GNN-DOVE selected a decoy within an iRMSD of 4 Å (one of the criteria for the acceptable quality level in CAPRI). This is in sharp contrast with the other methods (Figure 3A), where the iRMSD of many targets they selected was larger (worse) than GNN-DOVE. In terms of iRMSD, the second best method was DOVE, where 44 targets were within an iRMSD of 4 Å. A similar situation was observed for lRMSD. GNN-DOVE selected a decoy within a lRMSD of 10 Å (one of the criteria for the acceptable quality level in CAPRI) for 50 targets, while the second-best method, DOVE, selected 45 targets within 10 Å lRMSD. In terms of fnat (larger being more accurate), GNN-DOVE only missed 5 targets in selecting at least one model with an fnat over 0.1 (one of the criteria for acceptable quality level in CAPRI). The plot shows that GNN-DOVE had a larger fnat value than the other existing methods for most of the targets, as indicated by many data points below the diagonal line.

**Figure 3.**
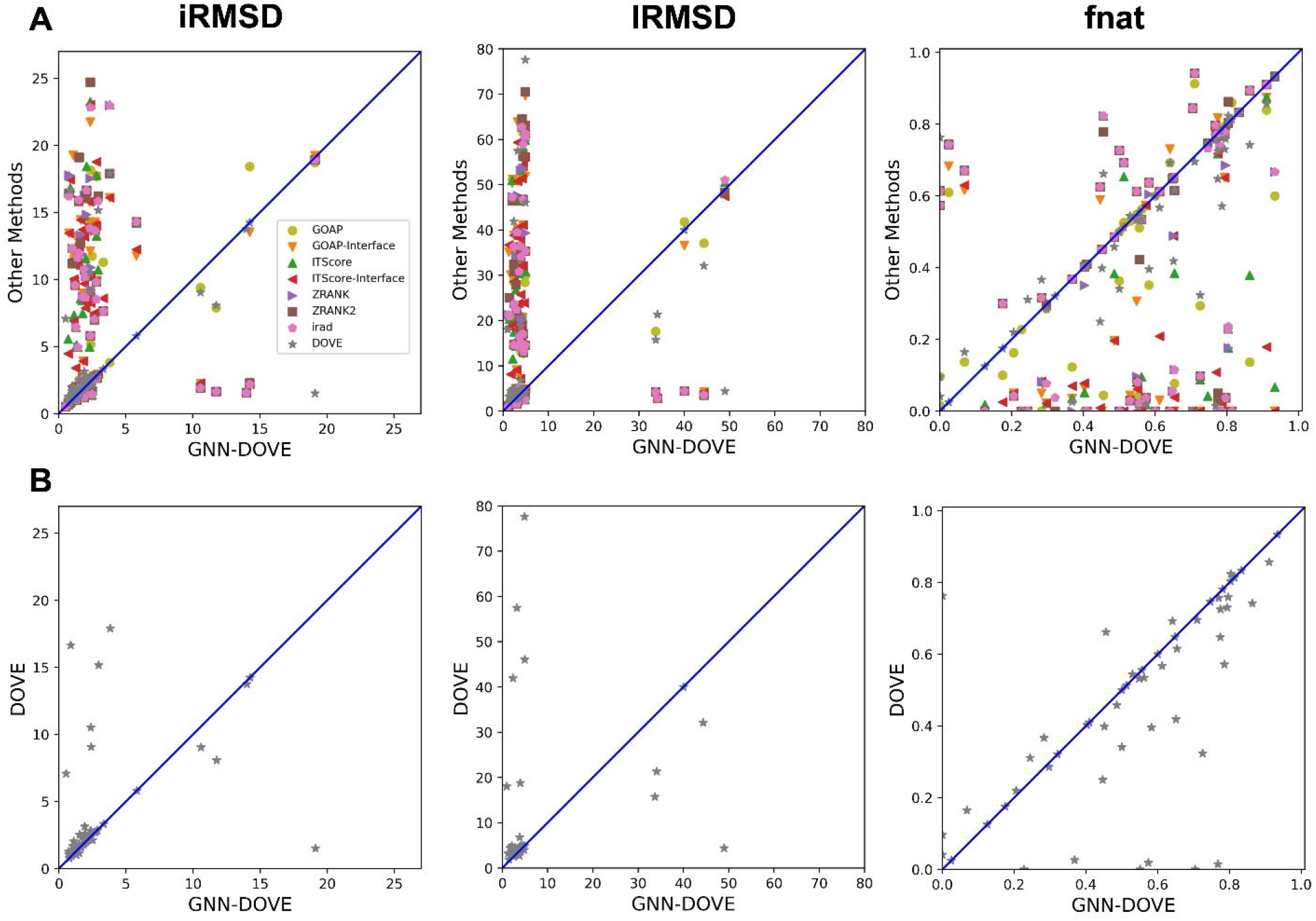
Comparison of iRMSD, lRMSD, and fnat. For each method, the best value among the top 10 scored decoys was plotted. **A**, Comparison against all eight methods. **B**, Comparison against DOVE.

Figure 3B compares GNN-DOVE against DOVE. In terms of iRMSD, lRMSD, and fnat, GNN-DOVE outperformed DOVE for 26 (22 ties), 27 (20 ties), and 27 (17 ties), respectively. Overall, GNN-DOVE outperforms the eight existing methods for all the three metrics.

### 3.2 T-SNE analysis

To illustrate how GNN-DOVE classified decoys, we used t-SNE (*54*) to visualize GNN-DOVE’s encoding of decoys in Figure 4. t-SNE is a dimension reduction method to visualize similarities of high-dimensional data points. Since we employed a four-fold cross-testing, a plot was provided for each of the four testing sets. In all the plots, particularly in Fold 3 and Fold 4, most of the acceptable decoys (black circles) were distinguished from incorrect ones (gray crosses), which indicates a good representation and generalization ability of the graph neural networks for this problem.

**Figure 4.**
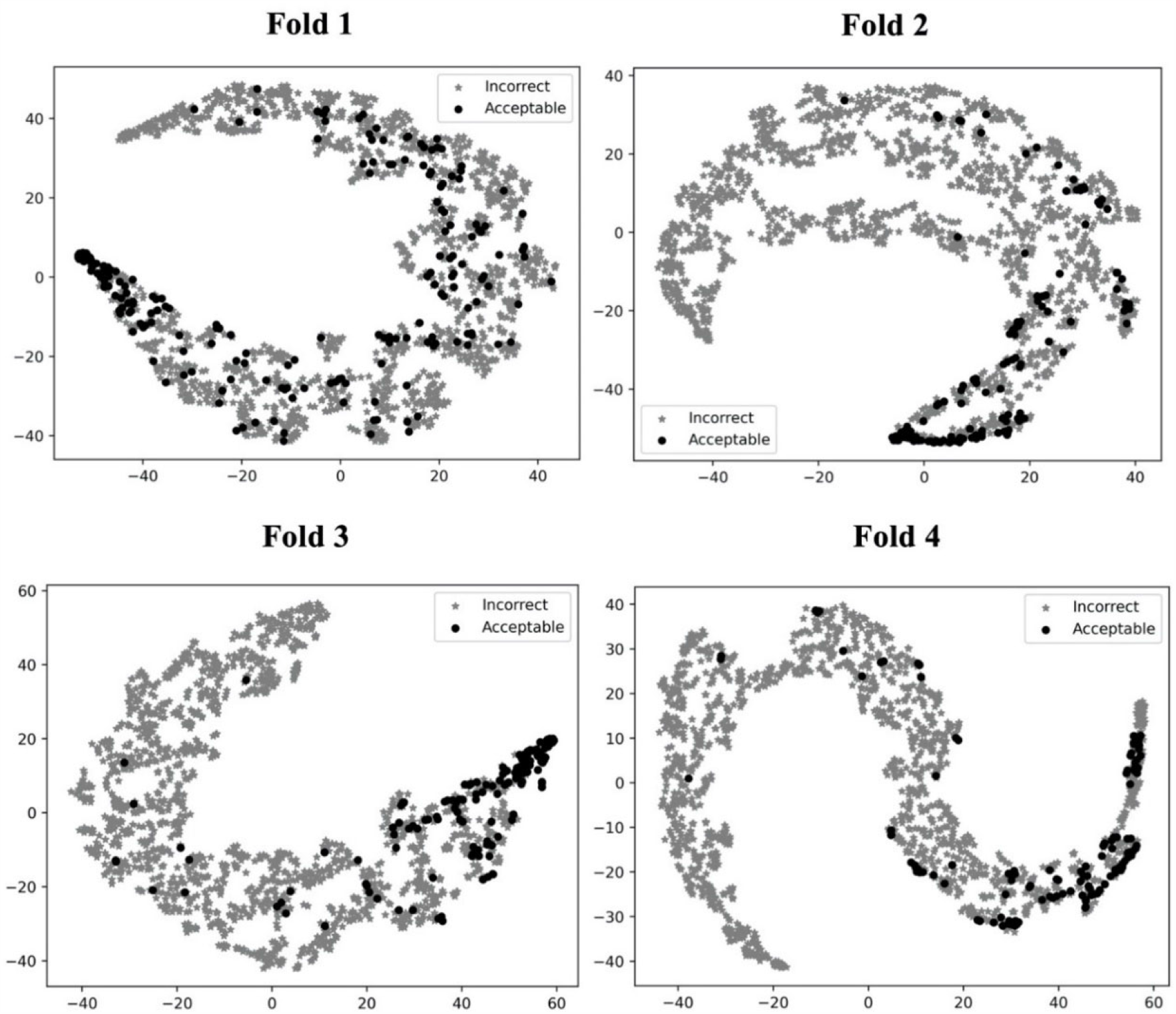
t-SNE plots of decoy selection. Decoys from all the testing target complexes in the four different folds in the cross-testing are plotted, which in total include 580 correct decoys (black circles) and 5591 incorrect decoys (gray stars). Encoded features of those decoys are taken from the output of the last fully connected layer of GNN, which is a vector of 128 elements. To visualize the different embedding, we use t-SNE to project them into a 2D space. The four panels correspond to the embedding of models on the four-fold testing sets.

### 3.3 Examples of decoys for comparison with DOVE

We mentioned above that a limitation of DOVE is that its usage of a fix-sized cube of 40^3^ Å^3^, which cannot capture the entire interface region if the interface is too large to fit in the cube. Here we show two examples of such cases, which led to a misclassification by DOVE but correctly classification by GNN-DOVE. In Figure 5, the interface region of a decoy is shown in blue and green, and the atoms that did not fit in the cube are shown in a sphere representation in red.

**Figure 5.**
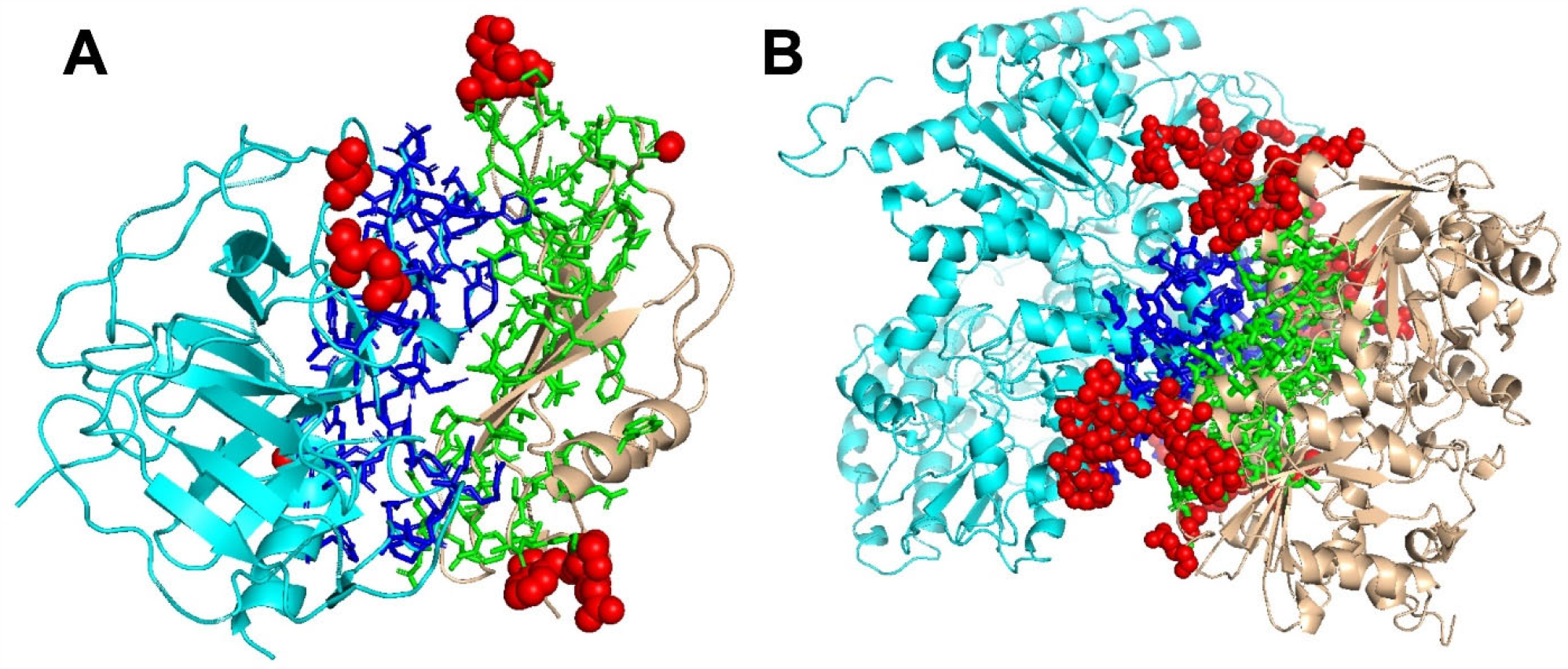
Examples of decoys with an acceptable quality but not selected within the top 10 by DOVE. Two subunits docked are shown in cyan and light brown and the interface regions of the two subunits are presented in the stick representation and in blue and green, respectively. To highlight the missed atoms from the input cube of DOVE, they are shown in red spheres. **A**, a medium quality decoy for 1bui. iRMSD: 2.54 Å; lRMSD: 2.93 Å; fnat: 0.551. **B**, a medium quality decoy for 1g20. iRMSD: 2.14 Å; lRMSD: 3.86 Å; fnat: 0.453.

The first example (Fig. 5A) shows a decoy of a protein complex of plasminogen and staphylokinase (PDB ID: 1bui), which has an acceptable quality by the CAPRI criteria. For this decoy, 59 atoms (in red) out of 1022 atoms at the interface were not included in the cube. Because of this, it was ranked the 65^th^ out of 110 decoys by DOVE, while it was ranked 15^th^ by GNN-DOVE. For this target, GNN-DOVE ranked five hits within the top 10 scoring decoys and eight hits within the top 20. In contrast, DOVE could not rank any hit within the top 20. The first hit by DOVE was found at the 35^th^ rank.

The second example (Fig. 5B) is an acceptable model for the nitrogenase complex (PDB ID: 1g20). As shown, many interface atoms, 497 out of 1843, were outside the cube. DOVE ranked this decoy 28^th^, while GNN-DOVE ranked this decoy 10^th^. DOVE had 0 hits within the top 10 and had only one hit within top 20. On the other hand, GNN-DOVE was very successful for this target, where all the top 10 selections were correct models.

## 4 Discussion

In this work, we developed GNN-DOVE for protein docking decoy selection, which used a Graph Neural Network (GNN). We used the gated augmented attention mechanism to capture the atom interaction pattern at the interface region of protein docking models. The benchmark on the Dockground dataset demonstrated that GNN-DOVE outperformed DOVE, along with other existing scoring functions compared.

To assess the quality of structure models, considering multi-body (atom or residue) interactions (*55-58*) have been proven to be an effective approach. GNNs consider patterns of multi-atom interactions by representing the interactions as a graph structure. Since a graph is a natural representation of molecular structures, GNNs may be applied in various problems in structural bioinformatics and cheminformatics.

The performance of GNN-DOVE likely would be improved by considering other physicochemical properties of atoms such as atom-wise binding energies, as well as sequence conservation of residues that can be computed from a multiple sequence alignment of homologous proteins. Application to multi-chain complexes remains a potential path for future work.

## Acknowledgements

The authors are grateful to Jacob Verburgt for proofreading the manuscript and Sai Raghavendra Maddhuri Venkata Subramaniya and Aashish Jain for testing the GNN-DOVE code on GitHub. This work was partly supported by the National Institutes of Health (R01GM133840, R01GM123055), and the National Science Foundation (DMS1614777, CMMI1825941, MCB1925643, DBI2003635).

## Authors Contributions

XW and STF conceived the initial version of the study. XW and DK designed this work in the current form. XW developed the codes in communication with STF. XW performed the computation and XW and DK analyzed the results. XW wrote the initial draft of the manuscript and DK critically edited it. All authors have read and approved the manuscript.

## Reference

1. T. Aderinwale, C. W. Christoffer, D. Sarkar, E. Alnabati, D. Kihara, Computational structure modeling for diverse categories of macromolecular interactions. Current Opinion in Structural Biology 64, 1–8 (2020).

2. I. Anishchenko, P. J. Kundrotas, A. V. Tuzikov, I. A. Vakser, Structural templates for comparative protein docking. Proteins: Structure, Function, and Bioinformatics 83, 1563–1570 (2015).

3. N. Tuncbag, A. Gursoy, R. Nussinov, O. Keskin, Predicting protein-protein interactions on a proteome scale by matching evolutionary and structural similarities at interfaces using PRISM. Nature Protocols 6, 1341 (2011).

4. V. Venkatraman, Y. D. Yang, L. Sael, D. Kihara, Protein-protein docking using region-based 3D Zernike descriptors. BMC Bioinformatics 10, 407 (2009).

5. B. G. Pierce, Y. Hourai, Z. Weng, Accelerating protein docking in ZDOCK using an advanced 3D convolution library. PloS One 6, e24657 (2011).

6. E. Katchalski-Katzir et al., Molecular surface recognition: determination of geometric fit between proteins and their ligands by correlation techniques. Proceedings of the National Academy of Sciences 89, 2195–2199 (1992).

7. D. Padhorny et al., Protein–protein docking by fast generalized Fourier transforms on 5D rotational manifolds. Proceedings of the National Academy of Sciences 113, E4286–E4293 (2016).

8. D. Fischer, S. L. Lin, H. L. Wolfson, R. Nussinov, A geometry-based suite of moleculardocking processes. Journal of Molecular Biology 248, 459–477 (1995).

9. I. H. Moal, P. A. Bates, SwarmDock and the use of normal modes in protein-protein docking. International Journal of Molecular Sciences 11, 3623–3648 (2010).

10. T. Oliwa, Y. Shen, cNMA: a framework of encounter complex-based normal mode analysis to model conformational changes in protein interactions. Bioinformatics 31, i151–i160 (2015).

11. J. J. Gray et al., Protein–protein docking with simultaneous optimization of rigid-body displacement and side-chain conformations. Journal of Molecular Biology 331, 281–299 (2003).

12. J. Esquivel-Rodríguez, Y. D. Yang, D. Kihara, Multi-LZerD: Multiple protein docking for asymmetric complexes. Proteins: Structure, Function, and Bioinformatics 80, 1818–1833 (2012).

13. D. W. Ritchie, S. Grudinin, Spherical polar Fourier assembly of protein complexes with arbitrary point group symmetry. Journal of Applied Crystallography 49, 158–167 (2016).

14. D. Schneidman-Duhovny, Y. Inbar, R. Nussinov, H. J. Wolfson, Geometry-based flexible and symmetric protein docking. Proteins: Structure, Function, and Bioinformatics 60, 224–231 (2005).

15. N. Alam et al., High-resolution global peptide-protein docking using fragments-based PIPER-FlexPepDock. PLoS Computational Biology 13, e1005905 (2017).

16. M. Kurcinski, M. Jamroz, M. Blaszczyk, A. Kolinski, S. Kmiecik, CABS-dock web server for the flexible docking of peptides to proteins without prior knowledge of the binding site. Nucleic Acids Research 43, W419–W424 (2015).

17. M. Kurcinski, A. Badaczewska-Dawid, M. Kolinski, A. Kolinski, S. Kmiecik, Flexible docking of peptides to proteins using CABS-dock. Protein Science 29, 211–222 (2020).

18. L. X. Peterson, A. Roy, C. Christoffer, G. Terashi, D. Kihara, Modeling disordered protein interactions from biophysical principles. PLoS Computational Biology 13, e1005485 (2017).

19. L. X. Peterson, W. H. Shin, H. Kim, D. Kihara, Improved performance in CAPRI round 37 using LZerD docking and template-based modeling with combined scoring functions. Proteins: Structure, Function, and Bioinformatics 86, 311–320 (2018).

20. L. X. Peterson et al., Modeling the assembly order of multimeric heteroprotein complexes. PLoS Computational Biology 14, e1005937 (2018).

21. J. Esquivel-Rodríguez, D. Kihara, Fitting multimeric protein complexes into electron microscopy maps using 3D Zernike descriptors. The Journal of Physical Chemistry B 116, 6854–6861 (2012).

22. G. C. van Zundert, A. S. Melquiond, A. M. Bonvin, Integrative modeling of biomolecular complexes: HADDOCKing with cryo-electron microscopy data. Structure 23, 949–960 (2015).

23. P. Gainza et al., Deciphering interaction fingerprints from protein molecular surfaces using geometric deep learning. Nature Methods 17, 184–192 (2020).

24. B. Akbal-Delibas, R. Farhoodi, M. Pomplun, N. Haspel, Accurate refinement of docked protein complexes using evolutionary information and deep learning. Journal of Bioinformatics and Computational Biology 14, 1642002 (2016).

25. M. T. Degiacomi, Coupling molecular dynamics and deep learning to mine protein conformational space. Structure 27, 1034-1040. e1033 (2019).

26. I. H. Moal, M. Torchala, P. A. Bates, J. Fernández-Recio, The scoring of poses in protein-protein docking: current capabilities and future directions. BMC Bioinformatics 14, 286 (2013).

27. M. F. Lensink et al., The challenge of modeling protein assemblies: the CASP12-CAPRI experiment. Proteins: Structure, Function, and Bioinformatics 86, 257–273 (2018).

28. L. J. Kingsley, J. Esquivel-Rodríguez, Y. Yang, D. Kihara, M. A. Lill, Ranking protein-protein docking results using steered molecular dynamics and potential of mean force calculations. Journal of computational chemistry 37, 1861–1865 (2016).

29. S. Y. Huang, X. Zou, An iterative knowledge-based scoring function for protein–protein recognition. Proteins: Structure, Function, and Bioinformatics 72, 557–579 (2008).

30. H. Lu, L. Lu, J. Skolnick, Development of unified statistical potentials describing protein-protein interactions. Biophysical Journal 84, 1895–1901 (2003).

31. F. Fink, J. Hochrein, V. Wolowski, R. Merkl, W. Gronwald, PROCOS: computational analysis of protein-protein complexes. Journal of computational chemistry 32, 2575–2586 (2011).

32. A. A. Nadaradjane, R. Guerois, J. Andreani, in Protein Complex Assembly. (Springer, 2018), pp. 429–447.

33. X. Wang, G. Terashi, C. W. Christoffer, M. Zhu, D. Kihara, Protein docking model evaluation by 3D deep convolutional neural networks. Bioinformatics 36, 2113–2118 (2019).

34. J. Janin et al., CAPRI: a critical assessment of predicted interactions. Proteins: Structure, Function, and Bioinformatics 52, 2–9 (2003).

35. F. Scarselli, M. Gori, A. C. Tsoi, M. Hagenbuchner, G. Monfardini, The graph neural network model. IEEE Transactions on Neural Networks 20, 61–80 (2008).

36. Z. Wu et al., A comprehensive survey on graph neural networks. IEEE Transactions on Neural Networks and Learning Systems, (2020).

37. D. K. Duvenaud et al., in Advances in Neural Information Processing Systems. (2015), pp. 2224–2232.

38. J. Lim et al., Predicting drug–target interaction using a novel graph neural network with 3D structure-embedded graph representation. Journal of Chemical Information and Modeling 59, 3981–3988 (2019).

39. J. S. Smith, O. Isayev, A. E. Roitberg, ANI-1: an extensible neural network potential with DFT accuracy at force field computational cost. Chemical Science 8, 3192–3203 (2017).

40. R. Zubatyuk, J. S. Smith, J. Leszczynski, O. Isayev, Accurate and transferable multitask prediction of chemical properties with an atoms-in-molecules neural network. Science Advances 5, eaav6490 (2019).

41. S. Liu, Y. Gao, I. A. Vakser, Dockground protein–protein docking decoy set. Bioinformatics 24, 2634–2635 (2008).

42. Y. Zhang, J. Skolnick, Scoring function for automated assessment of protein structure template quality. Proteins: Structure, Function, and Bioinformatics 57, 702–710 (2004).

43. W. Torng, R. B. Altman, Graph convolutional neural networks for predicting drug-target interactions. Journal of Chemical Information and Modeling 59, 4131–4149 (2019).

44. K. He, X. Zhang, S. Ren, J. Sun, paper presented at the Proceedings of the IEEE conference on Computer Vision and Pattern Recognition, 2016.

45. I. Goodfellow, Y. Bengio, A. Courville, Y. Bengio, Deep learning. (MIT press Cambridge, 2016), vol. 1.

46. D. P. Kingma, J. Ba, paper presented at the International Conference on Learning Representations, 2015.

47. N. Srivastava, G. Hinton, A. Krizhevsky, I. Sutskever, R. Salakhutdinov, Dropout: a simple way to prevent neural networks from overfitting. The Journal of Machine Learning Research 15, 1929–1958 (2014).

48. X. Glorot, Y. Bengio, in Proceedings of the thirteenth international conference on artificial intelligence and statistics. (2010), pp. 249–256.

49. H. Zhou, J. Skolnick, GOAP: a generalized orientation-dependent, all-atom statistical potential for protein structure prediction. Biophysical Journal 101, 2043–2052 (2011).

50. B. Pierce, Z. Weng, ZRANK: reranking protein docking predictions with an optimized energy function. Proteins: Structure, Function, and Bioinformatics 67, 1078–1086 (2007).

51. B. Pierce, Z. Weng, A combination of rescoring and refinement significantly improves protein docking performance. Proteins: Structure, Function, and Bioinformatics 72, 270–279 (2008).

52. T. Vreven, H. Hwang, Z. Weng, Integrating atom-based and residue-based scoring functions for protein–protein docking. Protein Science 20, 1576–1586 (2011).

53. X. Wang, G. Terashi, C. W. Christoffer, M. Zhu, D. Kihara, Protein docking model evaluation by 3D deep convolutional neural networks. Bioinformatics 36, 2113–2118 (2020).

54. L. v. d. Maaten, G. Hinton, Visualizing data using t-SNE. Journal of Machine Learning Research 9, 2579–2605 (2008).

55. H. Kim, D. Kihara, Protein structure prediction using residue-and fragment-environment potentials in CASP11. Proteins 84 Suppl 1, 105–117 (2016).

56. H. Kim, D. Kihara, Detecting local residue environment similarity for recognizing near-native structure models. Proteins 82, 3255–3272 (2014).

57. P. Gniewek, S. P. Leelananda, A. Kolinski, R. L. Jernigan, A. Kloczkowski, Multibody coarse-grained potentials for native structure recognition and quality assessment of protein models. Proteins 79, 1923–1929 (2011).

58. K. Olechnovic, C. Venclovas, VoroMQA: Assessment of protein structure quality using interatomic contact areas. Proteins 85, 1131–1145 (2017).

